# Diet Modifies Colonic Microbiota and CD4^+^ T Cell Repertoire to Trigger Flares in a Novel Model of Colitis Induced by IL-23

**DOI:** 10.1101/262634

**Authors:** Lili Chen, Zhengxiang He, Alina Cornelia Iuga, Sebastião N. Martins Filho, Jeremiah J. Faith, Jose C. Clemente, Madhura Deshpande, Anitha Jayaprakash, Jean-Frederic Colombel, Juan J. Lafaille, Ravi Sachidanandam, Glaucia C. Furtado, Sergio A. Lira

## Abstract

A wealth of experimental data points to immunological and environmental factors in the pathogenesis of inflammatory bowel disease (IBD). Here we study the role of IL-23, the microbiome, and the diet in the development of colitis. To promote IL-23 expression in vivo, we generated a mouse model in which IL-23 was conditionally expressed by CX_3_CR1^+^ myeloid cells, upon cyclic administration of tamoxifen in a specific diet (diet 2019). IL-23 expression induced an intestinal inflammatory disease that resembled ulcerative colitis in humans with cycles of acute disease and remission. The relapses were caused by the diet switch from the conventional diet used in our facility (diet 5053) to the diet 2019, and were not dependent on tamoxifen after the first cycle. The switch in the diet modified the microbiota, but did not alter the levels of IL-23. Colitis induction depended on the microbiota and required CD4 T lymphocytes. Colitis-inducing CD4^+^ T cells were found in the mesenteric lymph node and large intestine during remission and were able to trigger disease when transferred to lymphopenic mice, but only upon diet modification. The CD4 TCR repertoire in the diseased recipient *Rag^-/-^* mice had reduced diversity associated with the expansion of dominant T cell clones. These findings reveal a critical role for IL-23 in generation of a CD4^+^ T cell population in mice that is sensitive to a modification of intestinal bacterial flora subsequent to a dietary manipulation. Dietary changes occurring in the context of altered IL-23 expression may contribute to the onset and progression of IBD.

## INTRODUCTION

Crohn’s disease (CD) and ulcerative colitis (UC) are two distinct phenotypic patterns of inflammatory bowel disease (IBD) affecting over 1.5 million people in Europe and almost one million people in North American ^1^. At the turn of the 21st century, IBD has become a global disease with newly industrialized countries now facing rising incidence, analogous to trends seen in the western world during the latter part of the 20th century^2^. Inflammatory bowel disease is associated with morbidity, mortality, and substantial costs to the health-care system. Both CD and UC are characterized by periods of asymptomatic remission interrupted by episodes of symptomatic disease flares or exacerbations. While its exact cause is unknown, IBD seems to be due to a combination of genetic predisposition and environmental factors ^3-6^. Genome-wide association studies identified polymorphisms in several genes, including in the Interleukin-23 (IL-23) receptor (IL-23R)^7, 8^. IL-23 is a heterodimeric cytokine formed by the IL-23-specific p19 subunit and the IL-12p40 subunit. IL-23 interacts with cells that co-express the IL-23R subunit and the shared IL-12R β1 chain^9^. A number of cell types have been described as IL-23R positive and IL-23 responsive, including Th17 cells, gdT cells, macrophages, dendritic cells (DC), and innate lymphoid cells (ILC)^10, 11^. Several experimental models support a role for IL-23 in colitis. Loss-of-function studies show that IL-23p19 is essential for chronic colitis development in *IL-10^−/-^* spontaneous colitis model^12^, CD45RB^hi^CD4^+^ T-cell transfer models^13^, *Helicobacter hepaticus*-driven colitis^14^, anti-CD40-induced acute innate colitis model^11, 15^, and chemically induced colitis^16^. Furthermore, recent evidence implicates IL-23-responsive Th17 cells and ILCs in colitis^11, 15^, but the function of cytokines downstream of IL-23 (e.g. IL-17, IL-22) is still controversial. Other studies show that IL-23R expression by CD4^+^ T cells is required for disease development in murine T-cell transfer colitis model^17^ and that IL-23R expression by ILCs is important for colitis development in an anti-CD40 antibody-induced acute innate colitis model^15^. Although mounting evidence suggests that IL-23 is relevant in IBD pathogenesis, no direct demonstration that increased IL-23 signaling causes colitis exists to date.

Despite many advances, it is still not clear what environmental factors trigger development of IBD, nor it is known what factors cause flares of UC and CD. Besides genetic factors, smoking, diet, microbiota, and stress, appear to contribute to IBD development or aggravation^18^. Dietary changes have been proposed as a key factor in the increasing incidence of IBD in developing nations. Several large longitudinal studies have pointed to a lower risk of IBD among people who consume more fruits and vegetables, and a higher risk in people who consume less of these and more animal fats and sugar^19^. Consumption of specific foods has also been associated with CD and UC flares ^19^. In mice, the contribution of specific genes and the microbiota to colitis has been extensively analyzed. Mice bearing specific gene alterations develop colitis, but a significant number of them do not develop colitis when raised in germ-free conditions, suggesting a critical role for genes and the microbiota in promoting disease ^4, 20^. Other animal studies suggest a critical role of dietary components in the onset and severity of colitis ^21-23^. Yet, the development of relapsing-remitting disease models has not been reported. Because of this limitation it has been difficult to evaluate the contribution of diet and microbiota changes to disease initiation and progression.

To examine the contribution of IL-23, the microbiota and diet to development of colitis, we created a novel mouse model in which IL-23 is conditionally expressed by fractalkine chemokine receptor positive (CX_3_CR1^+^) cells. CX_3_CR1^+^ macrophages and DCs are the main cells expressing IL-23 in the gut upon exposure to bacterial antigens ^24, 25^. Our results show that CX_3_CR1^+^-derived IL-23 expression triggers development of a colitis that is dependent on the microbiota and the diet, with diet-driven cycles of active disease (relapse/flares) followed by remission. The development of colitis in this model is dependent on the generation of a CD4^+^ T cell response to the gut microbiota that is elicited by changes in the diet. Colitis-inducing CD4^+^ T cells are found in the mesenteric lymph nodes (mLN) and large intestine during remission and are able to trigger disease when transferred to lymphopenic mice, but only upon diet modification. Collectively, our experiments reveal a critical role for IL-23 in generation of a CD4^+^ T cell population that is sensitive to modification of intestinal bacterial flora subsequent to a specific dietary manipulation. The demonstration of a critical role for the diet in eliciting disease in genetically prone organism strongly supports the hypothesis of multi-causality in the etiopathogenesis of IBD.

## MATERIALS AND METHODS

### Mice

C57BL/6, *Rag1^-/-^*, *Tcrd^-/-^* and *muMt^-/-^* mice were purchased from The Jackson laboratory (Bar Harbor, ME). Mice were maintained under specific pathogen-free (SPF) conditions at the Icahn School of Medicine at Mount Sinai. The generation of IL-23 conditional knock-in mice (*R23* mice and *R23FR* mice) is detailed in Figure S1. All the germ-free (GF) mice were bred in-house and housed in standard flexible film isolators in our GF animal facility. All animal experiments in this study were approved by the Institutional Animal Care and Use Committee of Icahn School of Medicine at Mount Sinai, and were performed in accordance with the approved guidelines for animal experimentation at the Icahn School of Medicine at Mount Sinai.

### Diet Treatment

All mice were raised on the basal diet 5053, which was purchased from LabDiet (St. Louis, MO). The basal diet 2019 (TD. 160647) was purchased from Envigo (Madison, WI). Tamoxifen (500mg/kg) (Sigma) was added to the Envigo diet 2019 (TD. 130968). *R23FR* mice and control *FR* mice were fed with tamoxifen diet during the indicated times shown as Figure 1A. After each cycle of TAM treatment, animals were switched back to the basal diet 5053.

**Figure 1.**
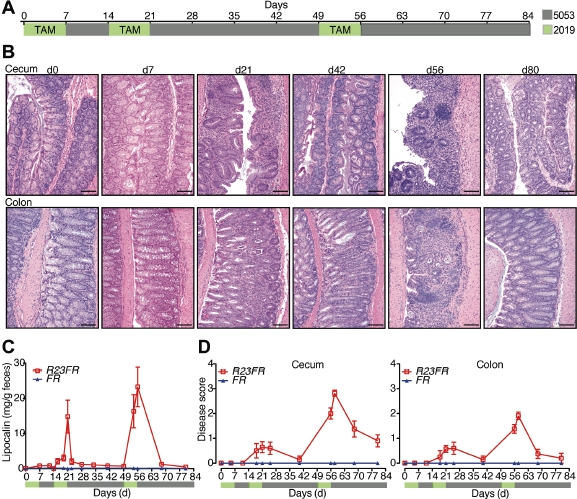
IL-23 Expression by CX_3_CR1^+^ Cells Induces Colonic and Cecal Inflammation. (*A*) Tamoxifen (TAM) in diet 2019 (green) was fed to *R23FR* and *FR* mice during the indicated times. After each cycle of TAM treatment, animals were switched to our mouse facility diet 5053 (gray). (*B*) Representative H&E-stained cecum and colon sections of *R23FR* mice at different time points. Scale bars, 100 μm. (*C*) Fecal lipocalin-2 levels in the stools of *R23FR* and *FR* mice were measured by ELISA (n= 5-15 per group per time point). (*D*) Histological scores of the colon and cecum of *R23FR* and *FR* mice at different time points (n= 5-15 per group per time point). Error bars represent mean ± SEM.

### Quantification of Fecal Lcn-2 by Enzyme-linked Immunosorbent Assay (ELISA)

Freshly collected or frozen fecal samples were reconstituted in PBS containing 0.1% Tween 20 (100 mg/ml) and vortexed for 20 min to get a homogenous fecal suspension^26^. These samples were then centrifuged for 10 min at 12,000 rpm and 4°C. Clear supernatants were collected and stored at −20°C until analysis. Lcn-2 levels were estimated in the supernatants using Duoset murine Lcn-2 ELISA kit (R&D Systems, Minneapolis, MN).

### Intraperitoneal Injection of Recombinant IL-23

*R23FR* mice after TAM treatment for 49 days were injected intraperitoneally (i.p.) with 2μg recombinant mouse IL-23 (R&D System) per mouse. Injections were continued in this manner every other day for 3 times (days 49, 51, and 53). Mice were sacrificed at d56, and gut was taken for histological analysis.

### Flow Cytometry and Sorting

Cell suspensions from the lamina propria and mLN were prepared as described previously^27^. All cells were first pre-incubated with anti-mouse CD16/CD32 for blockade of Fc γ receptors, then were washed and incubated for 40 min with the appropriate monoclonal antibody conjugates in a total volume of 200 μl PBS containing 2 mM EDTA and 2% (vol/vol) bovine serum. 4,6-diamidino-2-phenylindole (DAPI) (Invitrogen) was used to distinguish live cells from dead cells during cell analysis and sorting. Stained CD4^+^ T cells (DAPICD45^+^CD3^+^CD4^+^CD8^-^) were purified with a MoFlo Astrios cell sorter (DakoCytomation). Cells were > 95% pure after sorting. The sorted cells were used for TCR sequencing.

### T cell Purification

For CD4^+^ T-cell isolation, mLNs and large intestine were digested in collagenase as described previously ^27^. CD4^+^ T cells were enriched by positive immunoselection using CD4-(L3T4) microbeads (Miltenyi Biotec). The magnetic-activated cell sorting (MACS) purified CD4^+^ T cells were used as donor cells in adoptive transfer experiments.

### Histology

Tissues were dissected, fixed in 10% phosphate-buffered formalin, and then processed for paraffin sections. Five-micrometer sections were stained with hematoxylin and eosin (H&E) for histological analyses. All the sections were evaluated for a wide variety of histological features that included epithelial integrity, number of goblet cells (mucin production), stromal inflammation, crypt abscesses, erosion, and submucosal edema. Severity of disease was then classified based on a modified version of the Histologic Activity Index as described before^28^. Briefly, the disease score in the large intestine was calculated as follows: grade 0: absence of epithelial damage, focal stromal inflammation or regenerative changes; grade 1: crypt abscesses in less than 50% of the epithelium. Diffuse stromal inflammation and/or regenerative changes; grade 2: crypt abscesses in more than 50% of the epithelium and focal erosion or cryptic loss. Diffuse and accentuated crypt distortion with stromal inflammation; grade 3: Pan-colitis, diffuse erosion and ulcers.

### Microbiota Transplantation

Cecal extracts pooled from 3-5 donor SPF mice were suspended in PBS (2.5 ml per cecum) and were administered (0.1 ml per mouse) immediately to *R23FR* and *FR* germ-free mice that were generated in house ^29^. Transplanted mice were maintained in separate filter-top gnotobiotic cages and two weeks after were used for the further experiments^30^.

### DNA Extraction, 16S rDNA Amplification, and Multiplex Sequencing

DNA was obtained from feces of mice using a bead-beating protocol^31^. Briefly, mouse fecal pellets (∼50 mg), were re-suspended in a solution containing 700μL of extraction buffer [0.5% SDS, 0.5 mM EDTA, 20 mM Tris-Cl] and 200μL 0.1-mm diameter zirconia/silica beads. Cells were then mechanically disrupted using a bead beater (BioSpec Products, Bartlesville, OK; maximum setting for 5 min at room temperature), followed by extraction with QIAquick 96 PCR Purification Kit. Bacterial 16S rRNA genes were amplified using the primers as described in Caporaso et al^32^. Sample preparation and analysis of 16S rDNA sequence were done as previously described^33^. The 16S rDNA data were analyzed with MacQIIME 1.9.1. OTUs were picked with 97% sequence similarity and the sequences were aligned to the Greengenes closed reference set with a minimum sequence length of 150bp and a minimum percent identity of 75%^34, 35^.

### Antibiotic Treatment

For reduction of the intestinal microbiota, a ‘cocktail’ of antibiotics containing ampicillin 1g/L, metronidazole 1g/L, neomycin 1g/L, and vancomycin 0.5g/L (Sigma Aldrich) in the drinking water was provided to the mice through their drinking water. Antibiotic treatment was renewed every week.

### Reverse-transcription Polymerase Chain Reaction

Total RNA from tissues/sorted cells was extracted using the RNeasy mini/micro Kit (Qiagen) according to the manufacturer’s instructions. Complementary DNA (cDNA) was generated with Superscript III (Invitrogen). Quantitative PCR was performed using SYBR Green Dye (Roche) on the 7500 Real Time System (Applied Biosystems) machine. Thermal cycling conditions used were as follows: 50 °C for 2min and 95 °C for 10 min, 40 cycles of 95 °C for 15 s, 60 °C for 1min, followed by dissociation stage. Results were normalized to the housekeeping gene Ubiquitin. Relative expression levels were calculated as 2^(Ct(Ubiquitin)-Ct7^). Primers were designed using Primer3Plus^36^.

### In Vivo Antibody Treatment

To deplete CD4^+^ and CD8^+^ cells in mice, *FR* and *R23FR* mice treated with TAM were administered 100μg anti-CD4 (GK1.5, BioXcell) or anti-CD8 (53-5.8, BioXcell) or isotype antibodies (rat IgG2b, LTF-2; rat IgG1, TNP6A7; Both from BioXcell) intravenously at day 48, 49, 50, 52 and 54. Mice were sacrificed at day 56, and colon and cecum were taken for histological analysis.

### T cell Adoptive Transfer

One million CD4^+^ from mLN and/or large intestine enriched by using MACS-beads (Miltenyi Biotech) were transferred into *Rag1^-/-^* recipient mice by intravenous (i.v.) injection.

### Bulk Sequencing of TCRs

Briefly, cells were suspended in Trizol (Invitrogen) and total RNA was isolated. Using poly T beads, mRNA was isolated and fragmented briefly to generate 600-800bp fragments by using mRNA-seq kit (Bioo Scientific Corporation). Each of these RNA fragments were then converted to double stranded cDNA using random primers, end repaired and an A base added to each 3’end of the fragment. Two universal DNA sequences (A and B) (compatible with Illumina sequencing instruments) were ligated to each end of the fragment respectively. To enrich the library for T cell receptor transcripts, two sets of PCR reactions were performed. The first PCR reaction was done using a primer complementary to the Constant region and the universal sequence A. The second, nested PCR was performed using another constant region primer and universal sequence A. In the end Illumina compatible amplicons were generated with greater than 90% specificity to the T cell receptor transcripts.

The unique molecular indexes (UMID) in the sequenced reads were used to remove PCR duplicates and the reads were mapped to annotated V and J segments, which are compiled from IMGT (http://www.imgt.org), genome annotations from USUC (http://genome.ucsc.edu) and data from previous sequencing experiments performed in-house. All sequences that unambiguously mapped to V and J segments were translated to amino-acid sequences, in all three frames. Complementary determining region 3 (CDR3) sequences have well-defined boundaries, which were used to identify the CDR3 fragments in the translated open reading frames. Various categories (V-J pairs, CDR3 sequences, V-CDR3-J combinations) were tabulated from the data. Clonally expanded alpha and beta CDR3 were identified from the data, and the samples were clustered on the basis of the categories.

### Statistical Analysis

All statistical analyses were performed with GraphPad Prism 5 software. Differences between groups were analyzed with Student’s *t* tests or nonparametric Mann-Whitney test. Statistical tests are indicated throughout the Figure legends. Differences were considered significant when p < 0.05 (NS, not significant, * p < 0.05, **p < 0.01, ***p < 0.001), and levels of significance are specified throughout the Figure legends. Data are shown as mean values ± SEM throughout.

## RESULTS

### IL-23 Expression Induces An Inflammatory Disease that Resembles UC in Humans

Although IL-23 appears to be relevant in IBD pathogenesis both in human and experimental colitis model, there is no direct evidence that IL-23 expression can cause colitis in adult immuno-competent mice. To define the role of IL-23 in the intestinal inflammation, we engineered mice in which IL-23 expression could be induced by tamoxifen (TAM) in a subset of myeloid cells, known to express it in the gut (CX_3_CR1^+^ cells). This was accomplished by first generating Rosa26-lox-STOP-lox-IL-23 mice (*R23* mice) (Supplementary Figure 1A). The *R23* mice were subsequently mated to CX_3_CR1^CreER^ mice (*FR* mice) ^37^ to generate *R23FR* mice (Supplementary Figure 1A and B). As expected, TAM treatment promoted Cre-mediated excision of the STOP cassette and expression of IL-23 in CX_3_CR1^+^ intestinal cells of *R23FR* mice (Supplementary Figure 1C and D). To examine whether IL-23 expression would promote intestinal inflammation, we treated 60-70 days old *R23FR* and control *FR* mice with three cycles of TAM (500mg/kg in the diet 2019, Harlan Teklad). After each TAM cycle, animals were fed the conventional diet used in our mouse facility (diet 5053, Lab Diet) (Figure 1A). To monitor inflammation in a non-invasive way we measured the levels of Lipocalin 2 (Lcn-2) in the stools ^26^. The levels of fecal Lcn-2 varied during treatment of *R23FR* mice (Figure 1C), but not *FR* mice (Figure 1C), suggesting that expression of IL-23 induced inflammation. Histological evaluation of the gut was performed next, at the time points specified in Figure 1A. The small intestine of the *R23FR* mice appeared normal and did not show infiltrates at the time points examined. The cecum and colon of *R23FR* mice at d7 did not present inflammatory infiltrates (Figure 1B). However, by day 21 we observed the presence of leukocytic infiltrates in the mucosa of colon and cecum of *R23FR* mice (Figure 1B). In the colon, the presence of crypt abscesses was noted, but no erosions were observed (Figure 1B). By day 42, these infiltrates were very mild or absent. By day 56 (d56) the cecum and colon of *R23FR* mice showed a severe colitis, with marked leukocytic infiltrate, crypt loss, epithelial damage, and ulcerations (Figure 1B & D). The mucosa was expanded by the presence of a mixed lymphoplasmacytic and histiocytic infiltrates. No granulomata or transmural infiltrates were observed. The mucosal-based colitis with predominant cecal and colonic involvement was strikingly similar to human ulcerative colitis. By d80, the infiltrates were very mild or absent (Figure 1B). Histological analysis of *FR* treated mice did not show any abnormality at any time points (Supplementary Figure 2). Collectively, these data show that IL-23 expression drives development of an intestinal inflammatory disease that resembles ulcerative colitis in humans. Of note, cycles of acute disease and remission, observed in humans^38^, were also observed in this animal model.

### Diet Change Induces Development of Relapse

One of the striking aspects of this model was the development of relapse upon administration of a third cycle of TAM. To investigate the factors contributing to development of relapse we first investigated if the levels of transgenic IL-23 fluctuated in the intestine as function of TAM treatment. To do so, we performed qPCR analysis of the transgenic-specific 2A mRNA expression (Supplementary Figure 1A). As expected, the 2A mRNA expression levels increased at d7, compared with non-treated *R23FR* mice (d0) (Supplementary Figure 3). However, subsequent treatments (d56) did not increase them further (Supplementary Figure 3). We also probed if repeated injection of recombinant mouse IL-23 into *R23FR* mice starting at d49 for elicited disease (Supplementary Figure 4A). None of the mice developed disease at day 56(Supplementary Figure 4B and C). The results suggested that the relapses were not function of increased levels of IL-23. We also investigated if the relapse was due to the anti-estrogen receptor activities of TAM (Figure 2). Animals were treated with 2 cycles of TAM and subsequently treated with diet 2019 with TAM (Group 1), diet 5053 without TAM (Group 2), or diet 2019 without TAM (Group 3) (Figure 2A). As expected, animals in Group 1 had marked colitis (Figure 2B-2C), as shown in Figure 1. Animals in Group 2 did not develop colitis (Figure 2B-2C). Surprisingly, animals in Group 3, which did not receive TAM, but were fed with the diet in which TAM was dissolved (diet 2019), developed marked colitis (Figure 2B-2C). Diet 2019 promoted development of colitis independent of the levels of IL-23, as measured by the levels of 2A mRNA expression (Supplementary Figure 5). Subsequent experiments showed that development of colitis required a first cycle of TAM, as administration of diet 2019 to *R23FR* mice using the 3 cycles paradigm did not elicit disease (Supplementary Figure 6). Collectively, these results indicate that: 1) the induction of disease is dependent on the initial treatment of animals with TAM, 2) the relapses were not a function of increased IL-23 expression nor a function of estrogen receptor-blocking properties of TAM, and 3) the relapses were caused by the diet switch (from diet 5053 to diet 2019).

**Figure 2.**
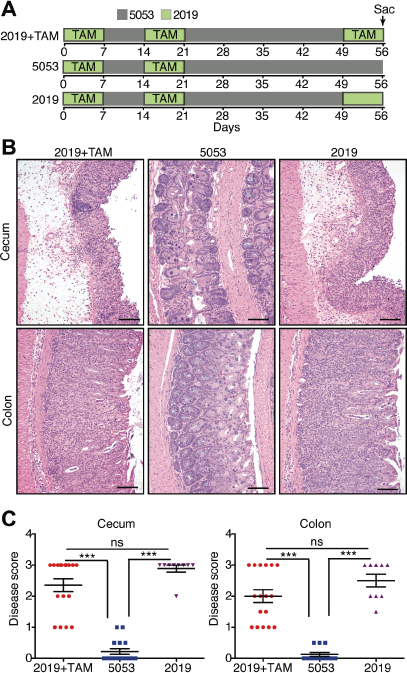
Change in Diet Causes Colitis Relapse in *R23FR* Mice. (*A*) Schematic representation of the diet switch experiments. (*B*) Representative H&E stained cecum and colon sections of *R23FR* mice at d56 from each group. Scale bars, 100 μm. (*C*) Histological scores of the colon and cecum of *R23FR* mice at d56. *** p<0.001; by nonparametric Mann-Whitney test.

#### Commensal Bacteria Are Required for IL-23 Induced Colitis

Many genetically susceptible animals do not develop colitis when raised in germ-free conditions ^4^. To test if microbes were required in our model, we developed a gnotobiotic colony of *R23FR* mice and *FR* mice, and treated them with the same protocol used for SPF mice (Fig 1). In striking contrast to the SPF *R23FR* mice, none of the GF *R23FR* mice showed inflammation in the cecum and colon at d56 (Supplementary Figure 7A). Next we asked if colonization of GF mice with commensal bacterial would be sufficient to cause disease. We found that introduction of C57BL/6 microbiota into GF *FR* mice did not cause disease, but that it caused colitis in GF *R23FR* mice (Supplementary Figure 7B). We then performed experiments to test whether different donor microbiota could induce disease (Supplementary Figure 8A). Transfer of SPF *R23FR* microbiota into GF *R23FR* recipient promoted disease, but this flora was not sufficient to cause disease when transferred into GF *FR* mice (Supplementary Figure 8B). No disease was observed in GF *FR* mice that received *FR* SPF microbiota (Supplementary Figure 8B). Transplant of the microbiota of SPF *R23FR* mice into *FR* mice did not cause disease suggesting that the *R23FR* microbiota at peak disease is not sufficient to induce disease and requires co-expression of IL-23. Together, these results indicate the microbiota is a key factor contributing to IL-23-induced colitis.

### Development of Relapse is Dependent on the Diet-modified Microbiota

Having established a critical requirement for the microbiota in disease development, we tested next if the induction of relapse induced by the diet switch required microbes. To do so, we treated *R23FR* mice and *FR* mice with 2 cycles of TAM and then with an antibiotic cocktail (ampicillin metronidazole, neomycin, vancomycin) from d35 to d56 (Figure 3A). All Antibiotic treatment significantly reduced colitis severity in the *R23FR* mice that received the diet 2019 from days 49-56 (Figure 3B-3C). These results further implicate the microbiota in the development of the diet-induced relapse.

**Figure 3.**
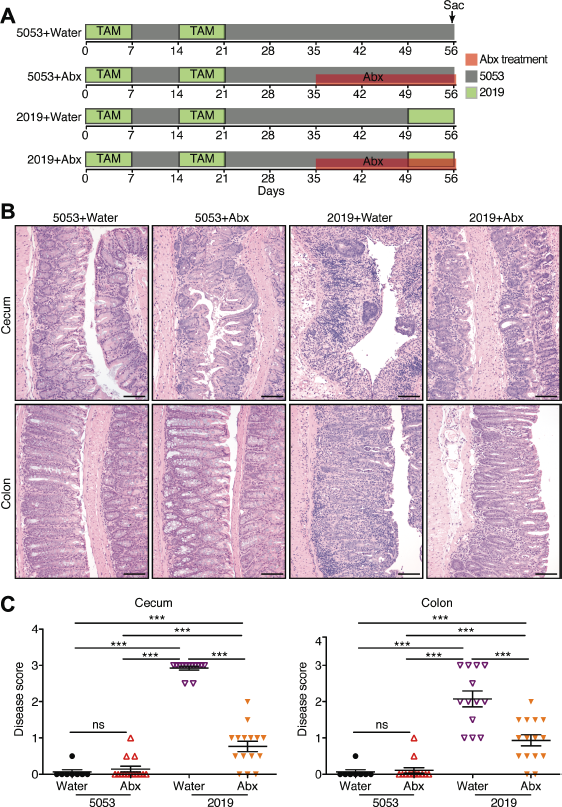
Microbiota is Required for Colitis Relapse. (*A*) Schematic representation of the experimental design. (*B*) Representative H&E stained sections of cecum and colon of *R23FR* mice at d56 from each group. Scale bars, 100 μm. (*C*) Histological scores of the colon and cecum of *R23FR* mice at d56. *** p<0.001; by nonparametric Mann-Whitney test.

To examine if the diet changes promoted changes in the composition of the microbiota we performed a longitudinal study with *R23FR* and *FR* mice. We collected fecal samples for microbiota analysis at different time points (d49, d51, d53 and d56, d58 and d63) (Figure 4A), and evaluated disease by histology (Figure 4B). Cecum inflammation was not observed in animals at d49. Diet switch promoted colitis in 2/11(∼18%) of the animals at d51, 5/11(∼46%) of the animals at d53 and 10/11 (∼91%) of the animals at d56 (Figure 4B). There was an increase not only in penetrance but also in severity of the disease during this time (Figure 4B). Analysis of the microbiome during this period revealed that there was a significant reduction in alpha diversity (a measure of overall microbiome diversity) in both groups as early as 4 days after the diet switch (d53). A very pronounced difference was observed between groups at d56 (Figure 4C). Further analysis showed that there were no significant taxa differences between *R23FR* and *FR* mice at d49 (prior to treatment), indicating that changes in diversity were not due to pre-treatment differences in the microbiome (Figure 4D). However, by d51, two days after diet switch, there was a significant expansion of *Verrucomicrobia* and a decrease of *Bacteroidetes* compared with day 49 in both *R23FR* and *FR* groups (Figure 4D and Supplementary Figure 9). Similar changes were found at d53 and d56 (Figure 4D and Supplementary Figure 9). At d56 there was a significant difference in the relative abundance of *Proteobacteria* in *R23FR* mice compared to *FR* mice, which coincided with peak inflammatory disease (Figure 4D and Supplementary Figure 9). Re-conversion to the 5053 diet (d58 and d63) resulted in rapid restoration of the microbiota to pre-treatment levels (d49) (Figure 4D). These results indicate that diet switch promoted significant changes in the microbiota of both *R23FR* and *FR* groups of mice at the time points examined (Figure 4D). Overall, our results indicate that development of flares caused by diet switch is dependent on the intestinal microbiota, and that a diet switch can rapidly change the composition of the microbiota.

**Figure 4.**
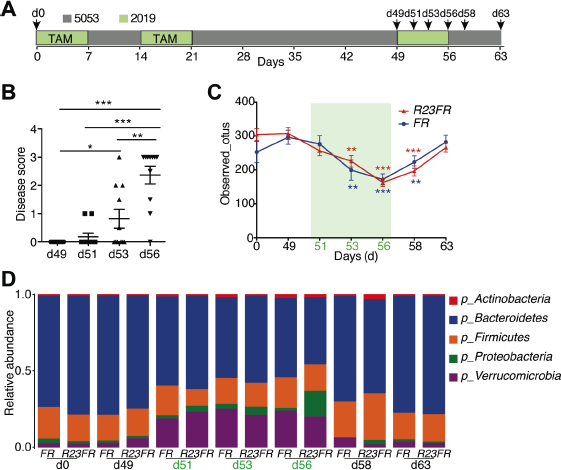
Diet Switch Changes the Composition of the Intestinal Microbiota. (*A*) Schematic representation depicts the experimental design. (*B*) Histological scores of the colon and cecum of *R23FR* mice at d49, d51, d53 and d56. *p<0.05, **p<0.01, *** p<0.001; by nonparametric Mann-Whitney test. (*C*) Alpha diversity (estimated as number of observed OTUs) at a sequencing depth of 12,000 seqs/sample. Asterisks (colored according to the different genotyping group) indicate a statistically significant difference in the number of observed OTUs compared to OTUs at day 49 within each group. **p<0.01, *** p<0.001; by nonparametric Mann-Whitney test. (*D*) Bacterial phylum-level community composition in the stool of *R23FR* and *FR* mice at each time point.

### Lymphocytes Are Required for Intestinal Inflammation

To examine the number and type of leukocytes present in the large intestine during disease we performed flow cytometry in cells obtained from the cecum of *R23FR* and *FR* mice. The overall number of leukocytes (as estimated by the number of CD45^+^ cells) present in the cecum of *R23FR* mice was not different from that of controls (*FR* mice) by the end of the first cycle of treatment (day 7) (Supplementary Figure 10A). The number of leukocytes rose significantly in the cecum of *R23FR* mice at the end of the second cycle (day 21), but fell afterwards. By day 42, the number of leukocytes was similar to that of *FR* mice. At the end of the third cycle, the number of leukocytes in the cecum of *R23FR* mice was, again, markedly increased (Supplementary Figure 10A). These results are consistent with the histological results presented before (Figure 1B & 1D). Next we performed a phenotypic analysis of the leukocyte subsets present in the large intestine lamina propria of *FR* and *R23FR* mice at day 56 (Supplementary Figure 10B and C). We found that the relative and absolute number of B cells, CD4^+^ and CD8^+^ T cells was dramatically increased in *R23FR* compared to *FR* mice (Supplementary Figure 10B and C) in overall analyses. These results indicate that expression of IL-23 induced markedly accumulation of B cells and T cells in the lamina propria.

Given the significant changes in the numbers of lymphocytes (Supplementary Figure 10B and C) observed in the LI of the *R23FR* mice and relevance of these cells to pathogenesis in experimental models, we tested next whether they were required for disease. To do so, we crossed *R23FR* mice with Rag deficient (*Rag^-/-^*) mice, which lack B and T cells, to generate *R23FR/Rag^-/-^* mice and subjected them to the protocol described in Figure 2A. Disease was completely absent in *R23FR/Rag^-/-^* mice, as determined by histological analysis of the large intestine (Figure 5A & 5B). These results indicated that lymphocytes were required for disease. To discriminate between the major lymphocyte populations causing disease, we crossed *R23FR* mice into *muMt^-/-^* mice, which lack B cells. By the end of third cycle of TAM treatment, we observed similar histological scores for *R23FR* and *R23FR/muMt^-/-^* mice (Figure 5A & 5B), suggesting that B cells were not required for disease.

**Figure 5.**
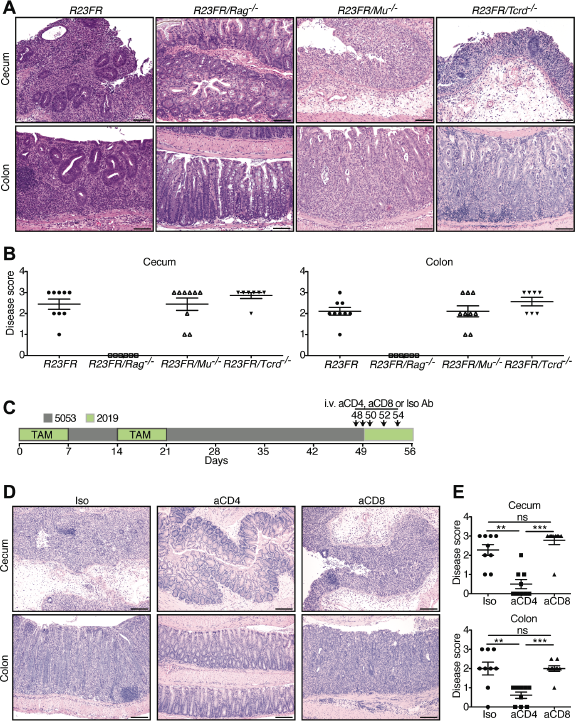
T Cells Are Required for Intestinal Inflammation in *R23FR* Mice. (*A*) Representative H&E stained sections from *R23FR, R23FR/Rag^-/-^, R23FR/Mu-/-* and *R23FR/Tcrd^-/-^* mice at d56. Scale bars, 100 μm. *(B) Histological scores of the colon and cecum of R23FR, R23FR/Rag^-/-^, R23FR/Mu^-/-^ and R23FR/Tcrd-/-* mice at d56. (*C*) Schematic representation depicts experimental design for cell depletion experiments. (*D*) Representative H&E-stained colon and cecum sections (left) and histological scores (right) of *R23FR* mice after treatment with anti-CD4 and anti-CD8 antibodies. **p<0.01, *** p<0.001; by nonparametric Mann-Whitney test.

### CD4^+^ T Cells Are the Main Effector Cells Promoting Colitis Upon Diet Switch

Since our results strongly suggested that T cells were the effector cells promoting colitis, we examined next which type of T cells accounted for the disease in the *R23FR* mice. To do so we crossed *R23FR* mice into *Tcrd^-/-^* mice that lack gamma-delta T cells. *R23FR/Tcrd^-/-^* mice had a disease that was indistinguishable from that observed in *R23FR/Tcrd^+/+^ mice* (Figure 5 & 5B), suggesting that TCRαβ T cells, not TCR gamma delta cells, were responsible for disease in the *R23FR* mice. To define which TCRαβ T cells were responsible for disease, we used antibodies to deplete CD4^+^ or CD8^+^ T cells (Figure 5C and Supplementary Figure 11). Depletion of CD4^+^, but not CD8^+^ T cells significantly reduced disease severity (Figure 5D) suggesting that CD4^+^ T cells had a critical effector role in this model.

To investigate if the CD4^+^ T cells obtained from *R23FR* mice could induce disease when transferred into *Rag^-/-^* mice (Figure 6A), we transferred MACS purified mLN and colonic CD4^+^ T cells from d49 *R23FR* and *FR* mice to *Rag^-/-^* mice and treated them with different diets for 21 days (Figure 6A). Transfer of control *FR* CD4^+^ T cells did not elicit colitis in *Rag^-/-^* mice, regardless of the diet regimen (Figure 6B). Transfer of *R23FR* CD4^+^ T cells to *Rag^-/-^* mice that were fed with diet 5053 also did not promote disease. However *Rag^-/-^* mice that received *R23FR* CD4^+^ T cells and were fed with 2 cycles of 2019 diet developed severe colitis (Figure 6B). Of note, there were comparable frequencies of CD4^+^ T cells in the blood of the recipients after adoptive transfer of CD4^+^ T cells (Supplementary Figure 12). Next we examined if both mLN and colonic CD4^+^ T cells could cause disease after transfer into *Rag^-/-^* mice. To do so, we purified *R23FR* d49 CD4^+^ T cells from mLN or large intestine separately and transferred them into *Rag^-/-^* mice (Figure 6C). Both mLN and colonic CD4^+^ T cells caused severe colitis in reconstituted *Rag^-/-^* mice after a diet switch (Figure 6D). These results indicate that CD4 T cells capable of inducing disease can be found in both mLN and colon of *R23FR* mice, but can only induce disease upon a diet-switch.

**Figure 6.**
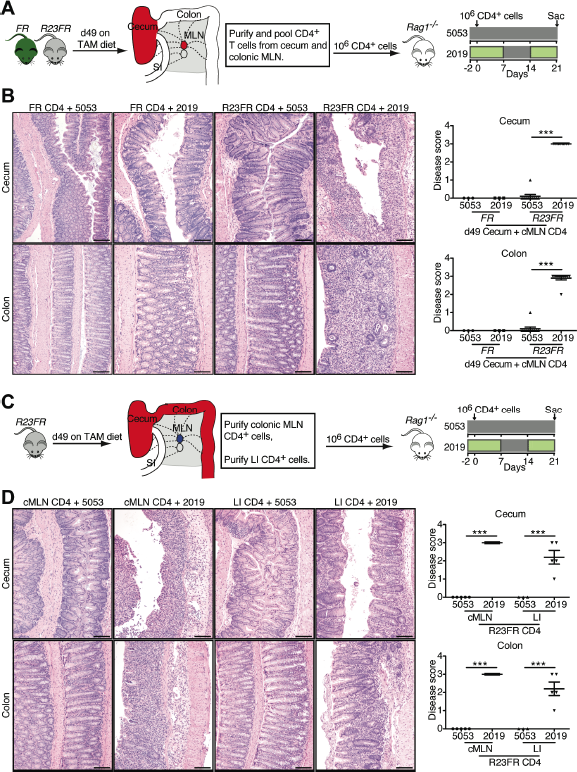
IL-23-induced CD4^+^ T Cells Drive Inflammation in Adoptively Transferred Mice. (*A*) Experimental setup for adoptive transfer of pooled CD4^+^ T cells from cecum and mLN of *R23FR* or *FR* mice into *Rag^-/-^* mice. (B) Representative H&E staining (left) and histological scores (right) of the colon and cecum of *Rag^-/-^* mice that received *R23FR* and *FR* mice CD4^+^ T cells fed with different diets. *** p<0.001; by nonparametric Mann-Whitney test. (C) Experimental setup for adoptive transfer of CD4^+^ T cells from large intestine or colonic mLN (cMLN) of *R23FR* mice into *Rag^-/-^* mice fed with different diets. (D) Representative H&E stained sections (left) and histological scores (right) of the colon and cecum of *Rag^-/-^* mice that received mLN or intestinal CD4^+^ T cells fed with different diets. *p<0.05, *** p<0.001; by nonparametric Mann-Whitney test.

### Differential Expansion of T cell Clones in Adoptively Transferred Mice Fed with Distinct Diets

The T cells in the gut specifically recognize antigens by virtue of their heterodimeric αβ T-cell receptor (TCR). Once T cells encounter their cognate antigen, they proliferate to produce clonal copies of themselves. In the homeostatic stage, the intestinal mucosal T cell repertoire is shaped by environmental antigens. Under chronic intestinal inflammation, the TCR repertoire is thought to be more profoundly shaped by immune response against lumen antigens^39^. We hypothesized that IL-23 expression resulted in generation of CD4^+^ T cells present in mLN and colon that could rapidly respond to microbiome changes induced by the diet. To examine if there was selective expansion of T cell clones upon diet switch, we sorted CD4^+^ T cells from donor *R23FR* mice, transferred them to recipient *Rag^-/-^* mice treated with different diets (Figure 7A), and sequenced both alpha and beta TCR chains from T cells recovered from the cecum at the end of treatment (Figure 7A). We clustered the sample populations on the basis of the complementary determining region 3 (CDR3) profiles, using hierarchical clustering. As expected, all donor cells from mLN for both alpha and beta chains clustered together (Figure 7B). Cells recovered from the cecum of three individual recipient *Rag^-/-^* mice treated with diet 2019 grouped together and apart from the cells recovered from recipient *Rag^-/-^* mice treated with diet 5053, and from mLN donor cells (Figure 7B). In addition, analysis of inverse Simpson’s diversity index revealed that the both alpha and beta chains of T cells from *Rag^-/-^* mice treated with 2019 diet had lower diversity compared to those of donor cells (Figure 7C). The decreased diversity of TCR repertoire in the cells from *Rag^-/-^* mice treated with diet 2019 was associated with expansion of T cell clones illustrated by an increase in the number of hyper-expanded clones (comprising >1% of all TCR sequences, Supplementary Table 1) of both TCRα and TCRβ (Figure 7D and Supplementary Table 2 and 3). These hyper-expanded clones present in the cecum of recipient *Rag^-/-^* mice treated with diet 2019 take up ∼25% of the total TCR repertoire, while they take up less than 10% in *Rag^-/-^* mice treated with 5053 diet (Figure 7D). More importantly, a few of the dominant clones expanded in all three individual recipient *Rag^-/-^* mice treated with 2019 diet (Figure 7E and Supplementary Table 2 and 3). These results indicate that the TCR repertoire in the diseased recipient *Rag^-/-^* mice had reduced diversity associated with the expansion of dominant T cell clones.

**Figure 7.**
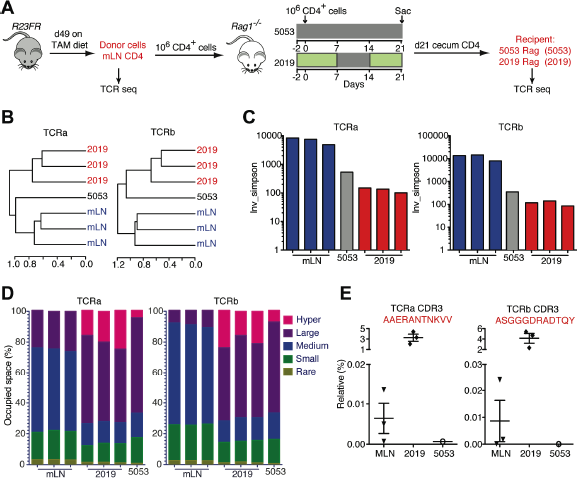
T Cell Receptor Clonality Analysis. (*A*) Experimental setup for TCR sequencing analysis. Note that donor cells from mLN of R23FR mice were obtained at remission stage (d49). (*B*) Unsupervised hierarchical clustering of TCR repertoire from donor mLN cells, cecum CD4+ T cells recovered from recipient Rag-/- mice treated with diets 5013 and 2019. N=3 individual mice in both donor group and 2019 group; n=1 of pooled 10 mice for group 5053. (*C*) Inverse Simpson’s diversity index was used to assess the diversity of TCR repertoires for each sample. (*D*) The percent of clonal space occupied by clones of a given type (classified by size, Supplementary Table 1). (*E*) Frequencies of selected TCRa and TCRβ clonotypes in different samples. primers F3 and R3 from untreated *R23FR* mice DNA, whereas a 500-bp fragment corresponding to the deleted allele was amplified from treated *R23FR* mice. (D) qPCR analysis of IL-23p19 and p40 mRNA expression from sorted YFP^+^ (CX_3_CR1^+^) cells from large intestine of *R23FR* mice without or with TAM treatment for 7 days. Columns and bars represent mean ± s.e.m (*p<0.5; **p<0.01, Mann-Whitney test).

## DISCUSSION

Here we show that low-level expression of IL-23 by CX_3_CR1^+^ cells triggered development of a colitis that was dependent on the microbiota and the diet, and that had a striking resemblance to human disease, with cycles of active disease (relapse or flares) followed by remission^38^. Expression of IL-23 resulted in the generation of a memory T cell population that was impacted by the microbiota, and had a skewed T cell repertoire. The colitogenic capacity of these T cells depended on the presence of a specific microbiota, which fluctuated in accordance to the diet.

IL-23 is a cytokine that has been implicated in the pathogenesis of several inflammatory conditions, including psoriasis, arthritis and IBD. Studies have shown that susceptibility to CD and UC is influenced by polymorphisms in the *Il23r* gene ^7, 8^, and that elevated levels of IL-23 are found in intestinal biopsies of patients with IBD ^40, 41^. Although IL-23 appears to be relevant in IBD pathogenesis both in human and experimental colitis models, there is no direct evidence to date that IL-23 expression can cause colitis in adult immunocompetent mice. Studies employing gene-modified mice have shown that mice expressing one of the IL-23 subunits (p19) in multiple tissues develop a multiorgan inflammatory disease that results in early lethality^42^. Dysregulated expression of IL-23 (both subunits) in the intestinal epithelium of transgenic mice induces severe intestinal inflammation in the small intestine, but not colon, and causes neonatal death^43^. Systemic exposure of adult animals to IL-23 via hydrodynamic delivery of plasmids encoding the IL-23 subunits results in rapid onset of psoriatic-like lesions in the ears, arthropathies in the paws and spine ^44, 45^, adenomatous tumors in the proximal duodenum ^46^, but not colitis. Here we show that IL-23 expression can lead to development of colitis; but that it requires simultaneous diet-driven changes in the microbiota to cause disease. These results add to the clinical evidence that targeting IL-23 through p40 (ustekinumab)^47^ or more specifically anti-p19 antibodies (brazikumab and risankizumab)^48, 49^ are effective in IBD.

The hypothesis that diet is an important factor contributing to the onset and progression of IBD, has remained formally unproven to date. Our results indicate that diet is a critical factor affecting development of IL-23-driven experimental IBD, and suggest that it does so by modifying the microbiota. Dietary constituents have been shown to affect the immune response and the inflammatory status, in part by modulating the microbiota ^22, 50^. Various dietary factors influence the growth of different bacterial populations and/or their functions in the gut. However, most studies to date have focused on how specific diet components (e.g. fat, carbohydrate, protein, fiber, et al.) promote or inhibit inflammation in experimental models ^50^. Analysis of the overall composition of the diets used in our experiments did not show significant differences in the content of fat, protein, fiber, or carbohydrates, so it is unclear what components are critical for the changes in the microbiota, but it is clear from our studies that colitis can not develop in the absence of the microbiota and that interference with the microbiota via antibiotic treatment prevents development of flares associated with the diet switch.

Our studies showed a marked reduction in alpha diversity as function of the diet change. These changes were already clear at d53 in both control and *R23FR* mice, and more pronounced at d56. Reduced diversity generally results in a reduced capacity of the microbial community to adapt to environmental pressures, and is likely to impact negatively on the microbial functional capacity ^51^. The taxa composition was similar between the groups at days 51 and 53, but was different from that of the same animals before the diet switch. The most evident change induced by the diet switch was an increased abundance of *Proteobacteria* and *Verrucomicrobia* in both groups. By d56, *R23FR* mice had an increased abundance of *Verrucomicrobia* and of *Proteobacteria* and a decrease in the abundance of *Bacteroidetes* compared to *FR* mice. The early increased abundance of *Verrucomicrobia* may have facilitated destruction of the mucus layer and adhesion of other bacterial species to the epithelium^52^. These changes, however, were not sufficient to promote colitis in *FR* mice, suggesting that the expression of IL-23 in the *R23FR* mice was critical for disease induction. IL-23 promotes expression of a number of cytokines that affect intestinal permeability^53^, the function of antigen-presenting cells^54^, regulatory T cells^55^ and memory cells ^56^.

The biological functions of IL-23 are mediated through the IL-23R, which is expressed by many cells of the innate and adaptive immune system, including Th17 cells, ILC3s ^11, 15^, gdT cells^57^ and neutrophils ^58^. Our studies showed that the colitis induced by IL-23 is dependent on lymphocytes, and that the other cell populations are not sufficient to drive disease. Interestingly, colitis in *R23FR* mice was unaffected by deficiency of B cells and gdT cells suggesting a critical role for CD3^+^ T cells. In addition, studies employing cell-depleting antibodies revealed a critical role for CD4^+^ T cells in pathogenesis, which was confirmed by adoptive transfer of these cells into *Rag^-/-^* mice. However, transfer of IL-23 induced CD4^+^ T cells was not sufficient to cause colitis. These cells could only elicit disease in response to an environmental modification (diet switch/microbiota changes). Thus, our findings provide the first evidence that IL-23 expression results in the generation of a colitogenic CD4^+^ T cell population induced by a dietary switch. In this context, we note that changes in the microbiota induced by diet have been hypothesized to contribute to the development of inflammatory bowel disease in humans ^59, 60^. Our results suggest that increased IL-23 signaling caused by intrinsic genetic defects or infection, may predispose the generation of pathogenic T cells. Indeed, the generation of long-term memory cells to commensals has been reported in mice infected with Toxoplasma^61^. The nature of the antigenic determinant eliciting the activation of the CD4^+^ T cell response in *R23FR* mice is not clear, but it appears to be derived from the bacteria, rather than from the diet. A food antigen would have triggered disease in the germ-free conditions, which was not observed here. However, factors other than bacterial antigens may also contribute to the overall immune response. As immune cells are sensitive to the action of bacterial metabolites, it is possible that in addition to bacterial antigens, they may be also responding to bacterial or host derived metabolites ^62^.

We propose that naive T cells become primed against bacterial antigens during the first cycle of TAM treatment. Expression of IL-23 and a concomitant change in the microbiota would result in generation of effector T cells. Effector T cells would migrate back to the intestine but would not respond with an inflammatory response since diet and microbiota would have reverted by then. These T cells would convert into memory cells and persist in the colon and mLN for extended periods of time, only to be activated upon diet-driven microbiota changes. The presence of microbiota-reactive T cells would not be detrimental on its own. Reactivity to intestinal bacteria is a normal property of the human CD4^+^ T cells present in the periphery and intestine^63^. Such auto reactive cells may occur as a function of previous pathogen exposure. Indeed, break in tolerance to commensals has been shown in mice previously exposed to infection with *T. gondii*^61^.

A fundamental property of memory T cells is the ability to proliferate rapidly and elicit effector functions in response to secondary challenge^61, 64^. We suggest that IL-23 expression favors development of a memory T cell population that can become pathogenic upon diet switch. As such, IL-23 could contribute to a break in tolerance to antigens present in the commensal microbiota. Indeed, a break in tolerance to commensal antigens is thought to induce chronic intestinal inflammation in humans^65^. Increased expansion and skewing of T cells has been reported in patients with IBD^39, 66^ and in the CD4^+^CD45RB^hi^ adoptive transfer mouse model^67^. The clonal expansion that we observed here in adoptive transferred *Rag^-/-^* mice fed with diet 2019 suggest that these expanded clones are pathogenic, but further studies are required to test this hypothesis.

In conclusion, we propose that IL-23 has a critical role in the generation of a CD4^+^ T cell population that reacts to bacterial shifts elicited by dietary manipulation. Our studies open new routes for the identification of dietary factors leading to expansion of the colitis-inducing microbiota and to the delineation of the immune mechanisms associated with the marked epithelial cell destruction and inflammation. The demonstration of a critical role for the diet in eliciting disease in genetically prone organism strongly supports the important role of diet, microbiome changes and IL-23 in the pathogenesis of IBD.

